# Interleukin-17 directly triggers Epstein-Barr virus lytic reactivation in latently infected human B cells

**DOI:** 10.64898/2025.12.19.695390

**Authors:** Janardhan Avilala, Hirotomo Dochi, Ramsy Abdelghani, David Becnel, Grace Maresh, Xin Zhang, Li Li, Mark Sides, Gilbert Morris, Hong Liu, Zhen Lin

**Author notes:** These authors contributed equally (Janardhan Avilala, Hirotomo Dochi).

## Abstract

Epstein-Barr virus (EBV) is a ubiquitous human gammaherpesvirus implicated in a wide spectrum of inflammatory and malignant diseases. Although EBV toggles between latent and lytic states, the host cues that control this switch remain incompletely defined. Interleukin-17A (IL-17A; hereafter IL-17), a signature Th17 cytokine, is abundant in EBV-associated tissues, and EBV products can augment IL-17 responses. Whether IL-17 directly modulates the EBV life cycle has remained unknown. Here we show that IL-17 alone is sufficient to induce EBV lytic reactivation in latently infected human B cells. Across RT-qPCR, immunoblotting, and RNA-seq, IL-17 increased the immediate-early regulator BZLF1 (Zta) and upregulated downstream early and late viral genes, consistent with activation of a transcriptome-wide lytic program. Culture supernatants from IL-17-treated cells contained elevated DNase-resistant extracellular EBV DNA, indicating productive replication with encapsidated genomes. Transcriptomic pathway analyses confirmed engagement of IL-17-linked signaling and highlighted inflammatory modules, including NF-κB and JAK-STAT. Gene Ontology analysis further enriched for regulation of B-cell receptor signaling and B-cell activation, situating IL-17 within B-cell inflammatory circuitry. Together, these findings identify IL-17 as a cytokine cue for EBV lytic entry and provide a mechanistic link between Th17-skewed inflammation and episodic EBV reactivation. This cytokine-driven pathway complements existing models centered on B-cell receptor signaling, hypoxia/HIF-1α, COX-2/PGE₂, and TGF-β and motivates testing whether modulation of the IL-17/IL-17R axis can alter EBV reactivation in IL-17-rich settings. To our knowledge, this is the first demonstration that IL-17 alone directly triggers EBV lytic reactivation in human cells.

**IMPORTANCE:** Epstein-Barr virus (EBV) persists for life by maintaining latency with intermittent lytic reactivation, yet the physiological cues that initiate the latency-lytic switch remain poorly defined. Here we identify the Th17 cytokine interleukin-17A (IL-17) as a direct host trigger of EBV lytic reactivation in latently infected human B cells. IL-17 alone induced the immediate-early transactivator Zta, activated a transcriptome-wide lytic program, and promoted release of DNase-resistant extracellular EBV DNA consistent with encapsidated virions. Transcriptomic analyses confirmed engagement of IL-17-linked inflammatory modules and implicated convergence on signaling nodes shared with canonical B-cell receptor pathways. These findings establish a mechanistic link between Th17-skewed inflammation and EBV reactivation burden, providing a framework to interrogate how IL-17-rich microenvironments influence EBV dissemination and immunopathology and to evaluate whether targeting the IL-17/IL-17R axis can modulate EBV reactivation in disease settings.

## INTRODUCTION

Epstein-Barr virus (EBV) is a ubiquitous human γ-herpesvirus that establishes life-long infection and contributes to a spectrum of inflammatory and malignant diseases (1–8). A defining feature of EBV biology is its ability to toggle between latency and lytic replication. Entry into the lytic cycle hinges on the induction of the viral immediate-early transactivators BZLF1 (Zta) and BRLF1 (Rta), which launch the downstream lytic transcriptional cascade and enable production of progeny virions (9–11). Understanding what cellular cues flip this latency/lytic switch remains central to explaining EBV pathogenesis in tissues where inflammation, hypoxia, or other stressors are prominent (12, 13).

Multiple host pathways can precipitate EBV reactivation *in vitro* and *in vivo* (13–17). Canonical inducers include B-cell receptor-driven signaling and cellular stress responses that converge on PKC, MAPK (ERK, JNK, p38), PI3K, and ATM networks to activate the BZLF1 promoter (Zp). Additional physiologic triggers have been described. Hypoxia/HIF-1α can bind Zp to drive BZLF1.

Inflammatory eicosanoids such as COX-2/PGE2 promote reactivation via EP receptor signaling. TGF-β induces lytic entry in epithelial and other EBV-infected cells. Collectively, these studies establish that EBV is responsive to host stress and inflammatory cues. However, the role of cytokines as direct lytic triggers is far less defined.

Interleukin-17A (IL-17A), often referred to simply as IL-17, is the prototypical Th17 cytokine and has potent context-dependent effects on stromal and epithelial cells (18, 19). In this study, all experiments were performed with recombinant human IL-17A and for brevity, we hereafter refer to IL-17A as IL-17. IL-17 signals through heterodimeric receptors that typically include IL-17RA with IL-17RC (or other subunits), recruiting the adaptor ACT1 and TRAF proteins to activate NF-κB, AP-1/MAPK, and post-transcriptional programs that stabilize target mRNAs. These features allow IL-17 to reshape tissue responses and, in some settings, to intersect with oncogenic and stress-response pathways, which could in principle influence EBV’s lytic switch (18, 19).

IL-17/Th17-skewed inflammation is detected across several EBV-associated contexts (20–22). In nasopharyngeal carcinoma (NPC), a tumor tightly linked to EBV, tumor tissues are enriched for Th17 cells, and IL-17 can promote NPC cell proliferation *in vitro*, underscoring the functional relevance of this axis in EBV-associated microenvironments. Conversely, classical Hodgkin lymphoma shows nuanced patterns where Th17 engagement can be stronger in EBV-negative disease, highlighting disease-specific heterogeneity. These observations nevertheless situate IL-17 as a plausible modulator in EBV-related tissues (20–22).

Notably, the directionality most often documented in the literature flows from EBV to IL-17 (23–25). That is, EBV nucleic acids and proteins augment IL-17 production and/or Th17 polarization through endosomal TLR pathways or other innate sensors. For example, purified EBV DNA enhances IL-17 via TLR-dependent mechanisms in mice, and the EBV dUTPase (a pathogen-associated molecular pattern) can elicit Th17-type cytokines. These findings imply that EBV reactivation or antigen release could fuel IL-17-rich milieus, but whether IL-17, in turn, directly regulates the EBV life cycle has remained an open question.

Cross-species precedent suggests such a link is plausible. In a murine γ-herpesvirus model (MHV68), IL-17 signaling supports viral reactivation and de novo lytic infection, and genetic or pharmacologic disruption of IL-17 pathways attenuates the establishment of chronic infection (26). This work positions IL-17 as a pro-viral host cue in γ-herpesvirus biology, yet direct evidence in human EBV-infected cells has been lacking.

Here we address this gap. We asked whether IL-17 alone is sufficient to trigger EBV lytic reactivation in latently infected human B cells. Using RT-qPCR, immunoblotting, a lytic reporter, DNase-resistant extracellular EBV DNA, and RNA-seq, we show that IL-17 induces BZLF1 and downstream early/late viral genes, consistent with productive lytic replication. Line-specific Hallmark GSEA demonstrates concordant enrichment of inflammatory modules (Inflammatory Response, TNFα signaling via NF-κB, IFN-γ Response, IL-2/STAT5), and a pooled GO Biological Process analysis recovers regulation of B-cell receptor signaling and B-cell activation, suggesting convergence of IL-17R outputs with BCR-distal effectors. Transcripts for IL-17RA/IL-17RC and TRAF3IP2/ACT1 are readily detectable in EBV(+) B cell lines, and IL-17 modulates ACT1 abundance in a line-dependent manner. Against the backdrop of established inducers (hypoxia/HIF-1α, COX-2/PGE₂, TGF-β, BCR-PKC-MAPK), our results identify a cytokine cue that directly flips the EBV latency-lytic switch in human B cells. This work reframes cytokine signaling as an upstream regulator of EBV’s life cycle and motivates mechanistic studies of receptor dependence and shared nodes (e.g., TRAF6/TAK1/IKK**/**MAPKs) as well as translational tests of IL-17/IL-17R blockade in IL-17-rich disease settings.

## MATERIALS AND METHODS

### Cell culture

The EBV-positive Akata cell line (type I latency) was established from an EBV-positive Burkitt’s lymphoma from a Japanese patient with a t(8:14) chromosome translocation (27). The EBV-positive Mutu (Mutu I) cell line (type I latency) was derived from an EBV-positive Burkitt’s lymphoma biopsy specimen from a Kenyan patient with a typical t(8:14) chromosome translocation (27). Rael is an EBV-positive Burkitt’s lymphoma cell line (type I latency). SNK6 is an EBV-positive NK/T-cell lymphoma-derived (NK-lineage) cell line (type II latency) Raji and Mutu III are EBV-positive Burkitt’s lymphoma cell lines (type III latency).

DG75 is an EBV-negative Burkitt’s lymphoma cell line. 293 is an EBV-negative human embryonic kidney cell line. All cell lines were maintained in either Dulbecco’s modified Eagle’s medium (DMEM; ThermoFisher Scientific, Cat# SH30243.02) for adherent cells, or RPMI 1640 (ThermoFisher Scientific, Cat# SH30027.02) for suspension cells, supplemented with 10% fetal bovine serum (FBS; ThermoFisher Scientific, Cat# 10437-028), penicillin, streptomycin, and glutamine. Cells were grown at 37°C in a humidified 5% CO_2_-containing atmosphere.

### IL-17 treatment and neutralization

Recombinant human IL-17A (carrier-free; Peprotech Cat# 200-17-100UG) was reconstituted in sterile PBS containing 0.1% BSA. For dose-response experiments, 1 × 10^6^ of cells were treated with IL-17 at 10 - 1000 ng/mL. Unless otherwise indicated, cells were harvested 24 h after cytokine addition for RT-qPCR and immunoblot analyses. Supernatants for extracellular viral DNA assays were collected at the time points specified in the figure legends. Based on these titrations, 500 ng/mL IL-17 produced maximal and reproducible induction and was used for subsequent experiments unless otherwise stated.

For neutralization assays, IL-17A was mixed with neutralizing anti-human IL-17A monoclonal antibody (R&D Systems, Cat# MAB31711-100) at the indicated concentrations and preincubated for 1 h at 37°C to allow immune-complex formation prior to addition to cells. Mutu I cells were treated with IL-17A (100 ng/ml) preincubated with antibody (20 µg/ml), and Akata cells were treated with IL-17A (50 ng/ml) preincubated with antibody (40 µg/ml). Antibody doses were chosen within the manufacturer’s recommended neutralization range.

### Transfection

Transient transfection experiments were performed by using either a modified version of the calcium phosphate precipitation procedure for adherent cells (28) or the Amaxa electroporation method for suspension cells (29). For the calcium phosphate precipitation procedure, 1 x 10^6^ cells were plated onto 100-mm-diameter tissue culture dishes. The following day, the medium was replaced with 8 ml of fresh supplemented DMEM; 4 hours later, 0.5 ml of 1x HEPES-buffered saline (0.5% HEPES, 0.8% NaCl, 0.1% dextrose, 0.01% anhydrous Na_2_HPO_4_, 0.37% KCl [pH 7.10]) was mixed with a total of 30 µg of plasmid DNA (effector plasmids were added in the amounts indicated in the figure legends, and carrier DNA [pUHD10] was added to make a total of 30 µg). 30 µl of 2.5 M CaCl_2_ was added, and samples were mixed immediately. Precipitates were allowed to form at room temperature for 20 minutes before being added dropwise to cells. Cells were incubated at 37°C with 5% CO_2_ for 16 hours before the medium was replaced with 10 ml of fresh supplemented DMEM. For the Amaxa electroporation method, cells were placed in antibiotic-free RPMI medium two days before electroporation. For each transfection, 2 x 10^6^ cells in 100 μl of Nucleofector Solution R (Mirus, Cat# MIR50118) were electroporated with a total of 5 μg of plasmid DNA (containing both effector and carrier DNA (pUHD10)) and transferred to a six-well plate with 1.5 ml of medium per well. After 24 hours, 1.5 ml of fresh RPMI medium was added to each well. Cells were incubated for another 24 hours before harvesting.

### RNA preparation

Whole cell RNA preparations were carried out using TRIzol reagent (Thermo Fisher, Cat# 15596) according to the vendor’s recommended protocol.

### RT-qPCR analysis

cDNA was synthesized from total RNA using iScript gDNA Clear cDNA Synthesis Kit (BioRad, Cat# 172-5035). qPCR analysis was performed using iQ SYBR Green Supermix (Bio-Rad, Cat# 170-8882) on a Bio-Rad CFX96 instrument as follows: 1 μl of cDNA and 1 μl of 10 μM primers were mixed with 10 μl of SYBR green supermix and 8 μl nuclease-free H2O to a 20 μl reaction volume. Polymerase was activated and cDNA was denatured at 95°C for 5 minutes. cDNA was then amplified for 40 cycles with 15 s denaturation at 95°C, 60s annealing/extension and plate reading at 60°C. Melting curve analysis was performed at temperatures from 60°C to 90°C with 0.5°C increment per 5 s. Expression fold changes were calculated using the comparative CT method (2-ΔΔCT).

### Western blot analysis

The cells were pelleted at desired time point and washed with 1X PBS and then the cells were immediately suspended in 100 μl of 1× Laemmli sample buffer and boiled for 15 min at 95°C to denature the genomic DNA. Protein concentrations of the whole-cell extracts were measured with the Bio-Rad protein assay kit according to the manufacturer’s instructions. Equal amounts of protein were subjected to SDS-polyacrylamide gel electrophoresis and transferred to PVDF membranes (Whatman). The blots were blocked for 60 min in Tris-buffered saline containing 5% nonfat powdered milk and 1% FBS and then incubated with the primary antibody (in blocking buffer) overnight at 4°C. The blots were washed three times with 1× TBST (140 mM NaCl, 3 mM KCl, 25 mM Tris-HCl [pH 7.4], 0.1% Tween 20) (each wash was carried out for approximately 15 min). To detect the chemiluminescence signals, the blots were then incubated with horseradish peroxidase-conjugated secondary antibody (Bio-Rad) in blocking buffer for 1 h at room temperature. Blots were washed as described above and analyzed with an enhanced chemiluminescence detection system (Perkin-Elmer) according to the manufacturer’s recommendations, and filters were exposed to either Fuji Super RX films (Fujifilm) or Kodak image station 4000 mm (Kodak, NY). To detect the infrared fluorescence signal, an Odyssey infrared imaging system (Li-Cor Biosciences) was used with a secondary antibody (IRDye 680RD goat anti-rabbit antibody; Li-Cor; Cat# 925-68071). The following primary antibodies were used for Western blot analysis: anti-Zta MAb (Santa Cruz; Cat# sc-53904), anti-BMRF1 MAb (Santa Cruz; Cat# SC-58121), anti-VCA-p18 (Thermo Cat# MAB8184), anti-GAPDH (Santa Cruz; Cat# SC-47724).

### Analysis of DNase-resistant EBV DNA

Cell-free, virion-associated EBV DNA was quantified from culture supernatants as DNase-resistant EBV DNA. Cells were pelleted (300 × g, 10 min, 4°C), and supernatants were clarified (2,000 × g, 10 min, 4°C) and passed through 0.45-µm syringe filters to remove residual cells and debris. Clarified supernatants were aliquoted and subjected to DNase treatment or mock treatment in parallel. For DNase digestion, Turbo DNase I (Thermo Fisher) was added to a final concentration of 20 U/mL in the supplied buffer (final 5 mM MgCl₂, 1 mM CaCl₂) and incubated 60 min at 37°C. Reactions were stopped by adding EDTA to 10 mM and heating 75°C for 10 min, followed by lysis with proteinase K (200 µg/mL) and 0.5% SDS for 30 min at 55°C. Viral DNA was extracted from 200 - 500 µL of treated or mock-treated supernatant using the QIAamp MinElute Virus Spin Kit (Qiagen) and eluted in 50 µL of nuclease-free water. EBV DNA was quantified by qPCR targeting EBV genomic noncoding sequences using validated primer sets. Absolute copy numbers were interpolated from plasmid standards or Namalwa genomic DNA (two EBV genomes/cell) and expressed as copies per mL of supernatant.

### Luciferase Reporter Assay

EBV-negative B cell lines or 293 cells were maintained in complete RPMI-1640 and DMEM, respectively. Cells (1.0-1.5 × 10⁶ per reaction) were transfected with reporter plasmids by nucleofection (Amaxa/Lonza; protocol #G-16). 1.0 μg Zp-luc reporter was used per reaction. Following a 24h recovery, cells were transferred to fresh medium and stimulated with recombinant human IL-17 (final 500 ng/mL) for 48 h. Cells were harvested in passive lysis buffer, and luciferase activities were measured using the Luciferase Reporter Assay System (Promega) on a plate luminometer following the manufacturer’s instructions. Data are presented as relative luciferase units (RLU) and expressed relative to the unstimulated Zp-luc control, set to 1.0.

### RNA-seq analysis

Total RNA was isolated from cells using the RNeasy kit (Qiagen) with on-column DNase I treatment and assessed on an Agilent 2100 Bioanalyzer. Only samples with RIN ≥ 8 were sequenced. Strand-specific, poly(A)-enriched libraries were sequenced on a DNBSEQ platform (BGI) to generate paired-end 100-bp reads (PE100). Raw sequencing reads were aligned to a reference genome containing a human genome (hg38; Genome Reference Consortium GRCH38) plus a modified Akata-EBV genome (Akata - NCBI accession number KC207813.1 (27)). The alignments were conducted using the Spliced Transcripts Alignment to a Reference (STAR) aligner version 2.5.3 (--clip5pNbases 6, default options) (30) and were subjected to visual inspection using the Integrative Genomics Viewer (IGV) genome browser (31). Transcript data from STAR were then analyzed using the RSEM software (version 1.3.0 (32)) for quantification of human and EBV gene expression. Signal maps (i.e., the total number of reads covering each nucleotide position) were generated using the IGV tools, and read coverage maps were visualized using the IGV genome browser (31). The EBSeq software (33) was utilized to call statistically differentially expressed genes using a false discovery rate (FDR) less than 0.05. GSEA-Preranked was run on EBSeq Fold change/statistic (positive = up in IL-17) using MSigDB Hallmark (v2025.1; symbols) with gene-set permutations (n=10,000), min/max set sizes 15/500, and Collapse/Remap = No. Significance was assessed by NES and FDR q-value. To conduct GO:BP enrichment (pooled), a pooled IL-17 signature (Akata+Mutu) was generated from the combined differential expression analysis (one *p*-value per gene; consistent direction required). Over-representation testing used a one-tailed hypergeometric/Fisher’s exact test against GO:BP (Human), with Benjamini–Hochberg FDR correction where indicated; the gene universe was the union of expressed genes in either line. Dot plots display -log10(*P*) (or FDR) and gene ratio (overlap/term size).

### Statistical analysis

RT-qPCR, luciferase reporter assays, and DNase-resistant viral DNA measurements were analyzed using GraphPad Prism. All statistical comparisons were performed using unpaired two-tailed Student’s *t*-test. Dose-response curves were fitted using a four-parameter logistic (4PL) model to derive EC_50_ values with 95% confidence intervals. qPCR data were log_2_-transformed prior to statistical testing. All tests were two-sided, and data are presented as mean ± SEM unless otherwise stated. Statistical significance was defined as *P* < 0.05.

### Database accession number

The RNA-seq sequence data have been deposited in the NIH Sequence Read Archive (SRA) database under accession number PRJNA1390600.

## RESULTS

### IL-17 induces EBV immediate-early transcription

To test whether IL-17 initiates EBV reactivation, EBV+ Akata and Mutu I cells were exposed to recombinant IL-17 (IL-17A) across 10 - 1000ng/mL, and BZLF1 (Zta) transcripts were quantified by RT-qPCR. Zta increased with dose, first detectable at 10 - 100ng/mL and reaching a plateau at 500ng/mL (**Fig. 1A**).

**FIG 1.**
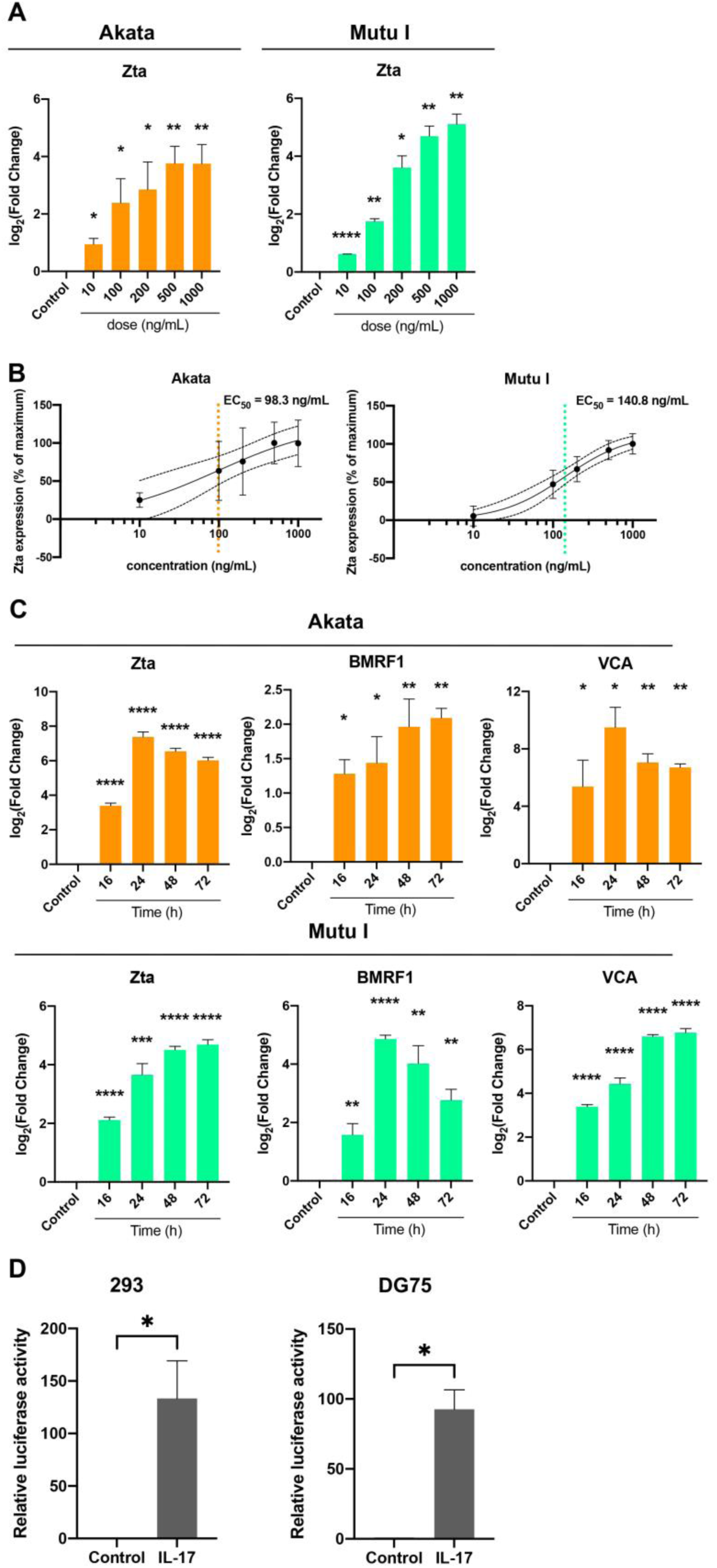
IL-17 induces EBV immediate-early transcription. (A) Akata and Mutu I cells were treated with increasing concentrations of IL-17 (0 to 1,000 ng/ml) for 48 h, and BZLF1 (Zta) mRNA was quantified by RT-qPCR. Expression was normalized to GAPDH and is shown as log_2_(fold change; 2⁻ΔΔCᴛ) relative to untreated cells. (B) Dose-response curves for Zta induction in Akata and Mutu I cells. Fold changes from panel A were fit by nonlinear regression (four-parameter logistic), yielding EC₅₀ values of ∼100 ng/ml (Akata) and ∼140 ng/ml (Mutu I). (C) Akata and Mutu I cells were treated with IL-17 (500 ng/ml), and mRNA levels of BZLF1 (Zta), BMRF1, and BLLF1 (VCA) were quantified by RT-qPCR at the indicated times (16 to 72 h). Expression was normalized to GAPDH and is shown as log_2_(fold change; 2⁻ΔΔCᴛ) relative to time-matched controls. (D) EBV-negative 293 and DG75 cells were transfected with a luciferase reporter driven by the BZLF1 promoter (Zp) and then treated with vehicle or IL-17 (500 ng/ml) for 24 h. Relative luciferase activity is shown normalized to control. For all RT-qPCR panels in this figure, bars/points represent means ± SEM from ≥ 2 biological replicates; each biological replicate was assayed in technical triplicate. Statistical significance was assessed by t test versus the corresponding control. *, P < 0.05; **, P < 0.01; ***, P < 0.001; ****, P < 0.0001.

Nonlinear regression of the Zta dose response (four-parameter logistic on log₁₀[IL-17]) yielded an EC_50_ of 98.3 ng/mL (95%CI, 28.7-228.7; Hill slope = 0.63) in Akata cells and 140.8 ng/mL (95%CI, 98.6-190.0; Hill slope =1.08) in Mutu I cells (**Fig. 1B**). Maximal induction was achieved at ≥500ng/mL.

We therefore used 500ng/mL for subsequent experiments to achieve maximal, reproducible activation. Although this dose exceeds circulating IL-17 levels, supraphysiologic cytokine concentrations are routinely used *in vitro* to model high local paracrine exposure, compensate for receptor abundance/affinity in cell lines, and offset cytokine instability. Similar ranges (100 - 500ng/mL) have been widely used to elicit robust signaling in human keratinocytes, fibroblasts, endothelial/platelet systems, and immune-cell assays (34–40).

At 500 ng/ml IL-17, Zta transcripts were already strongly induced by 16h in both Akata and Mutu I cells and remained elevated throughout the 72h time course (**Fig. 1C**). The early lytic gene BMRF1 and the late gene VCA also showed distinct temporal patterns of induction consistent with their positions in the lytic cascade. To determine whether IL-17 acts at the BZLF1 promoter (Zp), EBV-negative DG75 and 293 cells were transfected with a Zp-luciferase reporter. IL-17 increased Zp-driven luciferase activity relative to vehicle, consistent with promoter-level activation (**Fig. 1D**).

### IL-17 drives full EBV lytic reactivation and release of DNase-resistant viral DNA

We next asked whether IL-17 advances beyond immediate-early induction to a full lytic program. EBV+ Akata and Mutu I cells were treated with 500ng/mL IL-17 for 48h and analyzed by immunoblot and RT-qPCR. IL-17 increased immediate-early (Zta), early (EAD/BMRF1), and late (VCA) proteins, with concordant changes in lytic transcripts (**Fig. 2A-B**).

**FIG 2.**
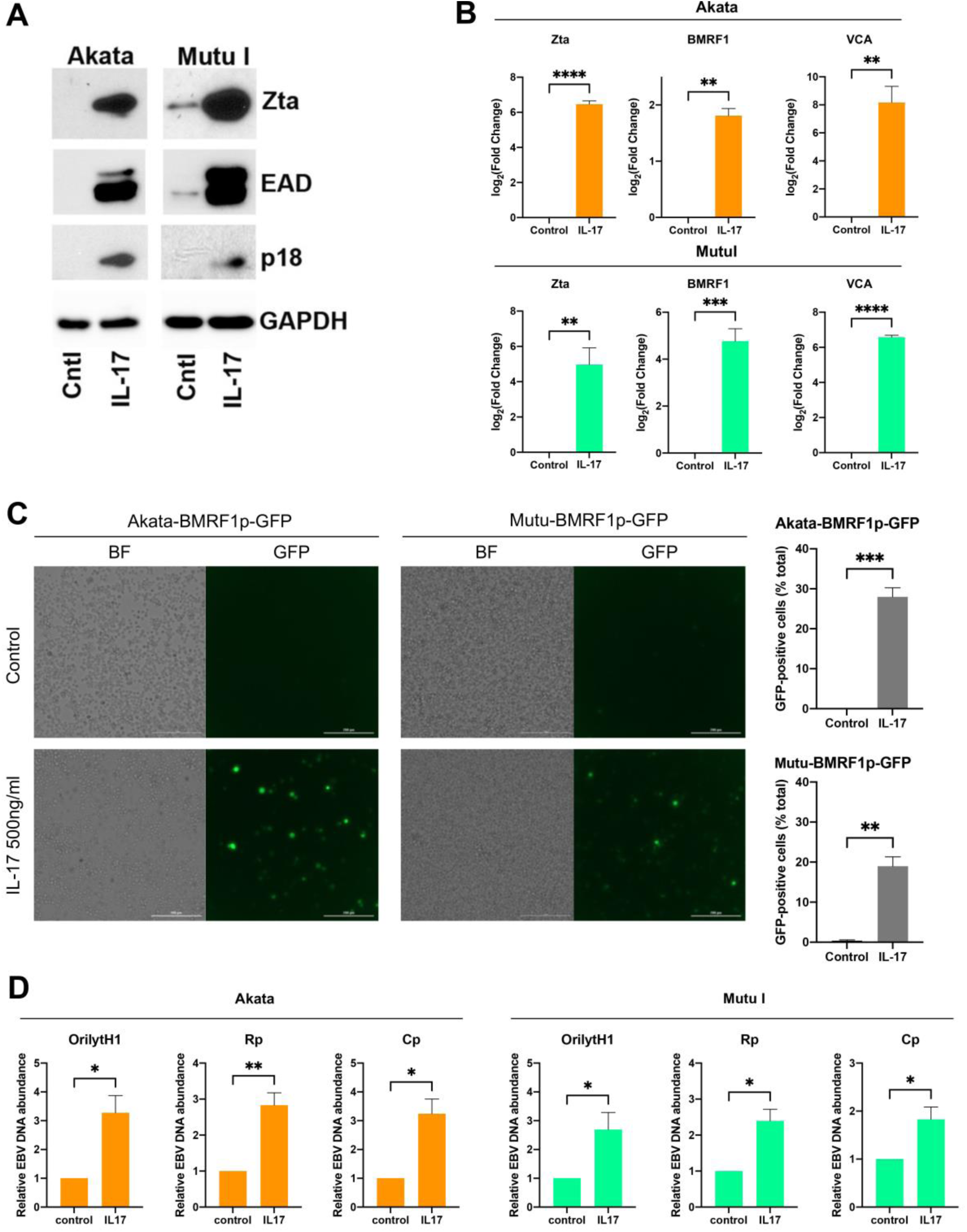
IL-17 drives full EBV lytic reactivation and release of DNase-resistant viral DNA. (A) Akata and Mutu I cells were treated with vehicle or IL-17 (500 ng/ml) for 48 h, and lysates were analyzed by immunoblotting with antibodies against Zta, EAD, and p18 (VCA). GAPDH served as a loading control. (B) RT-qPCR analysis of BZLF1, BMRF1, and BLLF1 (VCA) transcripts in Akata and Mutu I cells treated as in panel A. Expression was normalized to GAPDH and shown as fold change relative to control. (C) Akata and Mutu I cells stably expressing a lytic reporter (GFP under the BMRF1 promoter) were treated with vehicle or IL-17 (500 ng/ml) for 48 h. GFP-positive cells were quantified from fluorescence images using ImageJ and expressed as the percentage of GFP-positive cells. Scale bar: 200 μm. (D) DNase-resistant cell-free EBV DNA in culture supernatants from Akata and Mutu I cells treated with vehicle or IL-17 (500 ng/ml) for 72 h was quantified by qPCR. EBV genome copies were measured using three primer pairs targeting distinct noncoding/non-transcribed regions of the viral genome (Orilyt-H1, Rp, and Cp). For all RT-qPCR and qPCR panels in this figure (B and D), data represent means ± SEM from ≥ 2 biological replicates; each biological replicate was assayed in technical triplicate. Statistical significance was assessed by t test versus the corresponding control. *, P < 0.05; **, P < 0.01; ***, P < 0.001; ****, P < 0.0001.

An independent lytic reporter-GFP under the BMRF1 promoter was activated upon IL-17 treatment, indicating progression through the early lytic phase (**Fig. 2C**). Quantification of GFP-positive cells by immunofluorescence showed that approximately 28% of Akata cells and 19% of Mutu cells entered the lytic program after IL-17 treatment (48h) (**Fig. 2C**), which exceeds the ∼12-15% reactivation typically observed with anti-Ig stimulation in the same reporter system. These data support a population-level increase in lytic entry rather than a purely per-cell change in expression.

To determine whether IL-17 promotes productive events, we quantified DNase-resistant, cell-free EBV DNA in culture supernatants. IL-17 significantly increased nuclease-protected viral DNA in both Akata and Mutu supernatants (**Fig. 2D**), consistent with the presence of encapsidated genomes rather than naked DNA from dying cells. Together, these data indicate that IL-17 induces full lytic reactivation culminating in release of DNase-resistant viral DNA.

### IL-17-dependent reactivation is specific and neutralized by anti-IL-17

To establish ligand specificity, IL-17 was pre-incubated with a neutralizing anti-IL-17 antibody prior to addition to EBV+ cells. Furthermore, cells were also treated with heat-inactivated IL-17 as a control. We show that both neutralization and heat inactivation attenuated IL-17-induced lytic readouts including Zta/BMRF1/VCA transcripts (**Fig. 3**). These results demonstrate that the reactivation we observe is IL-17-dependent rather than a nonspecific response to protein addition.

**FIG 3.**
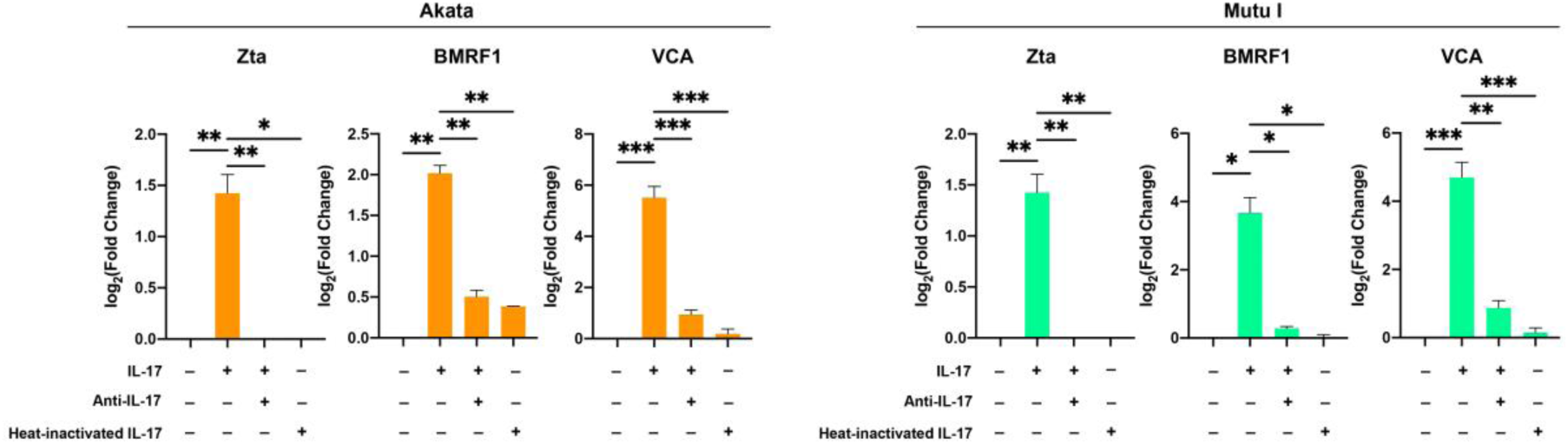
IL-17-dependent reactivation is specific and neutralized by anti-IL-17. Akata and Mutu I cells were treated with vehicle, IL-17, IL-17 preincubated with an anti-IL-17 neutralizing antibody, or heat-inactivated IL-17 as indicated. After 48 h, RT-qPCR was performed for EBV lytic transcripts (BZLF1, BRLF1, and BMRF1). Expression was normalized to GAPDH and shown as fold change relative to vehicle-treated cells. Bars represent means ± SEM from ≥ 2 biological replicates; each biological replicate was assayed in technical triplicate. Statistical significance was assessed by t test. *, P < 0.05; **, P < 0.01; ***, P < 0.001; ****, P < 0.0001.

### Transcriptome profiling links IL-17 signaling to EBV lytic gene expression and B-cell inflammatory programs

To define global transcriptional responses, we performed RNA-seq on EBV+ Akata and Mutu cells treated with IL-17 or vehicle. Reads were mapped to a combined human/EBV reference. Genome-wide visualization across the EBV genome revealed broad increases in immediate-early (BZLF1/BRLF1), early (e.g., BMRF1), and late (e.g., VCA/BLLF1) transcripts after IL-17 (**Fig. 4A**), consistent with full lytic progression and concordant with our DNase-resistant virion DNA measurements. Host volcano plots highlighted top 20 significantly altered cellular genes, with fewer repressed loci based on FDR values (**Fig. 4B**).

**FIG 4.**
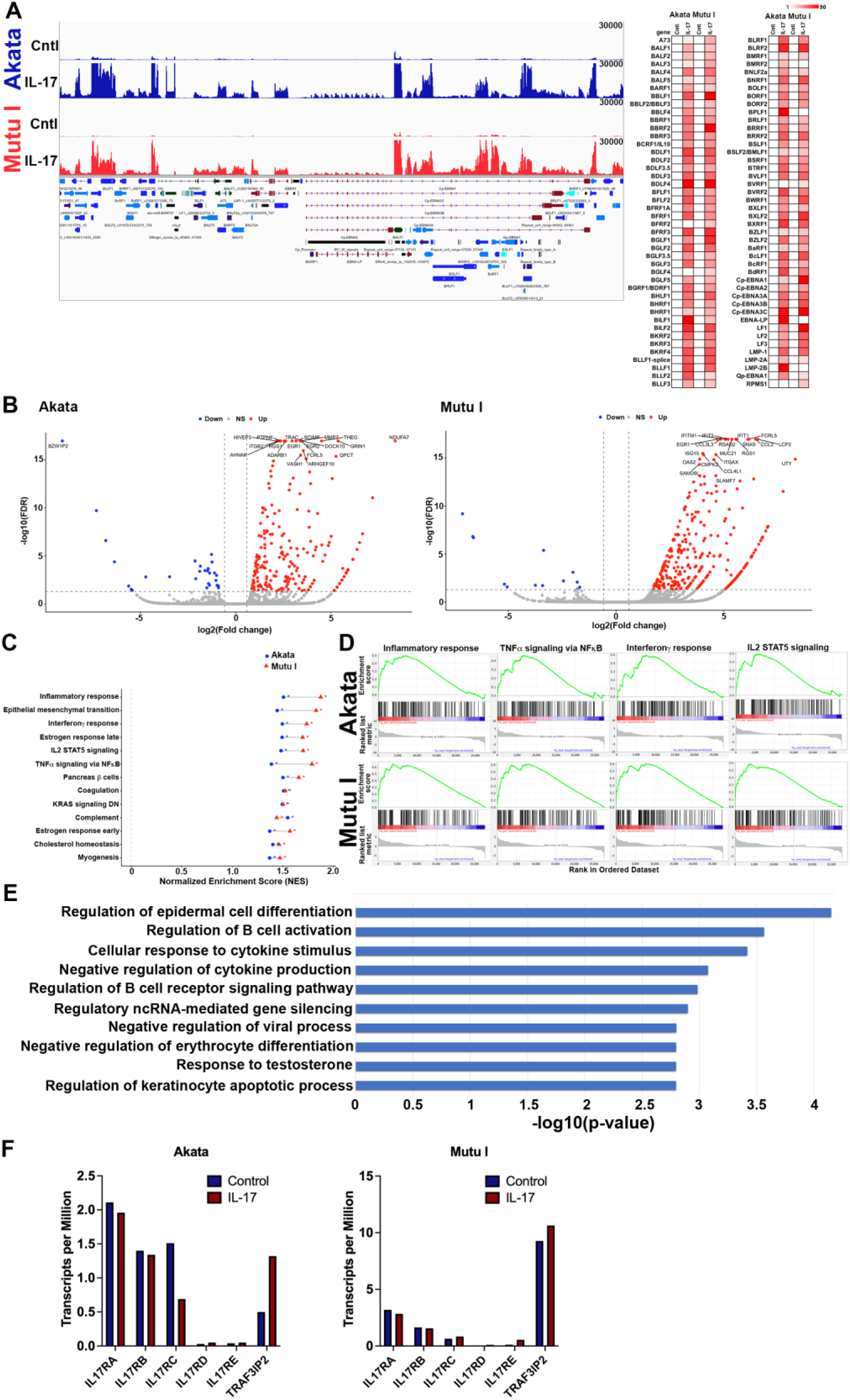
RNA-seq profiling corroborates lytic reactivation and engagement of IL-17 signaling. (A) Akata and Mutu I cells were treated with vehicle or IL-17 (500 ng/ml) for 48 h and subjected to DNBSEQ RNA-seq (paired-end 100 bp). Representative coverage across the EBV genome is shown using Integrative Genomics Viewer (IGV), together with a heat map of viral gene expression. (B) Volcano plots of host gene expression changes in Akata and Mutu I (IL-17 versus control). Dotted lines indicate thresholds (|log₂ fold change| ≥ log₂1.5; -log₁₀FDR ≥ 1.3). Selected top altered genes are labeled. (C and D) Hallmark GSEA-Preranked identified 13 pathways significantly enriched (FDR q < 0.05) in both Akata and Mutu I cells after IL-17 treatment. Normalized enrichment scores (NES) for shared pathways are shown. (E) Pooled Gene Ontology Biological Process over-representation analysis of IL-17-responsive genes from Akata and Mutu I highlights terms related to B-cell receptor signaling and B-cell activation. (F) RNA-seq-based TPM values for IL-17 receptor subunits (IL17RA, IL17RC) and the adaptor TRAF3IP2 (ACT1) in Akata and Mutu I cells treated with vehicle or IL-17.

To assess the alteration of host biological processes and signaling pathways, we next conducted the GSEA analysis using the human Hallmark gene sets as references. We identified 13 concordant pathways with FDR q<0.05 in both Akata and Mutu I cell lines (**Fig. 4C-D**). Enriched sets were dominated by inflammatory modules including inflammatory response, TNFα signaling via NFkB, interferon gamma response, and IL2-STAT5 signaling with positive NES in Akata (1.37-1.55) and higher NES in Mutu (1.44-1.88). Additional concordant terms include epithelial mesenchymal transition, complement, coagulation, cholesterol homeostasis, estrogen response early/late, myogenesis, pancreas beta cells, as well as KRAS signaling DN (i.e., Genes down-regulated by KRAS activation).

To increase functional resolution, we conducted an orthogonal GO Biological Process over-representation analysis on altered gene signatures after IL-17 treatment. The pooled GO analysis recovered B-cell-centric programs, including Regulation of B-cell receptor signaling pathway (GO:0050855; *P*≈1.0×10⁻³), Regulation of B-cell activation (GO:0050864; *P*≈2.7×10⁻^4^), and Cellular response to cytokine stimulus (GO:0071345; *P*≈3.8×10⁻^4^), together with Negative regulation of cytokine production (GO:0001818; *P*≈8.4×10⁻^4^), Negative regulation of viral process (GO:0048525; *P*≈1.6×10⁻³), and Regulatory ncRNA-mediated gene silencing (GO:0031047; *P*≈1.3×10⁻³) (**Fig. 4E**). These orthogonal analyses converge on IL-17-driven NF-κB/JAK-STAT activation with BCR/antigen-receptor-linked biology, aligning with our functional readouts of EBV lytic entry.

RNA-seq TPMs documented IL17RA and IL17RC, the canonical IL-17 receptor subunits in both EBV+ B-cell lines (**Fig. 4F**). IL17RA was readily detectable and minimally altered by IL-17. IL17RC was lower in abundance and changed in opposite directions. Other paralogs (IL17RB/D/E) were low to very low and showed small absolute changes. The adaptor TRAF3IP2 (ACT1) was present in both lines, with a ∼2.6-fold increase in Akata and a ∼15% increase in Mutu.

Together, these data indicate that an IL-17RA-IL-17RC/ACT1 signaling axis is available in both models. Short-term stimulation evokes modest, line-divergent transcript adjustments while preserving receptor/adaptor capacity consistent with the robust functional readouts of IL-17 signaling and EBV lytic induction.

### IL-17 preferentially promotes lytic entry in EBV latency I B-cell models

Because Akata and Mutu carry latency I, we asked whether IL-17 could also trigger reactivation in other EBV latency programs. EBV+ Rael (latency I), SNK6 (latency II), as well as Mutu III and Raji (latency III) were treated with 500 ng/mL IL-17 for 48 h, and assayed by western blot for Zta/EAD/VCA under identical conditions. Rael showed robust induction of reactivation with increased lytic proteins, mirroring the response in Akata/Mutu (**Fig. 5**). In contrast, SNK6, Mutu III and Raji exhibited no detectable increase in lytic proteins relative to vehicle controls (**Fig. 5**). These data indicate that IL-17-driven lytic entry is context-dependent and appears selective for latency I models. IL-17 alone was insufficient to trigger reactivation in the latency II or III lines tested. The latency-program specificity suggests that cell-state-dependent factors (e.g., promoter accessibility or cofactor availability) rather than receptor presence alone likely determine IL-17 responsiveness and will be probed in future work.

**FIG 5.**
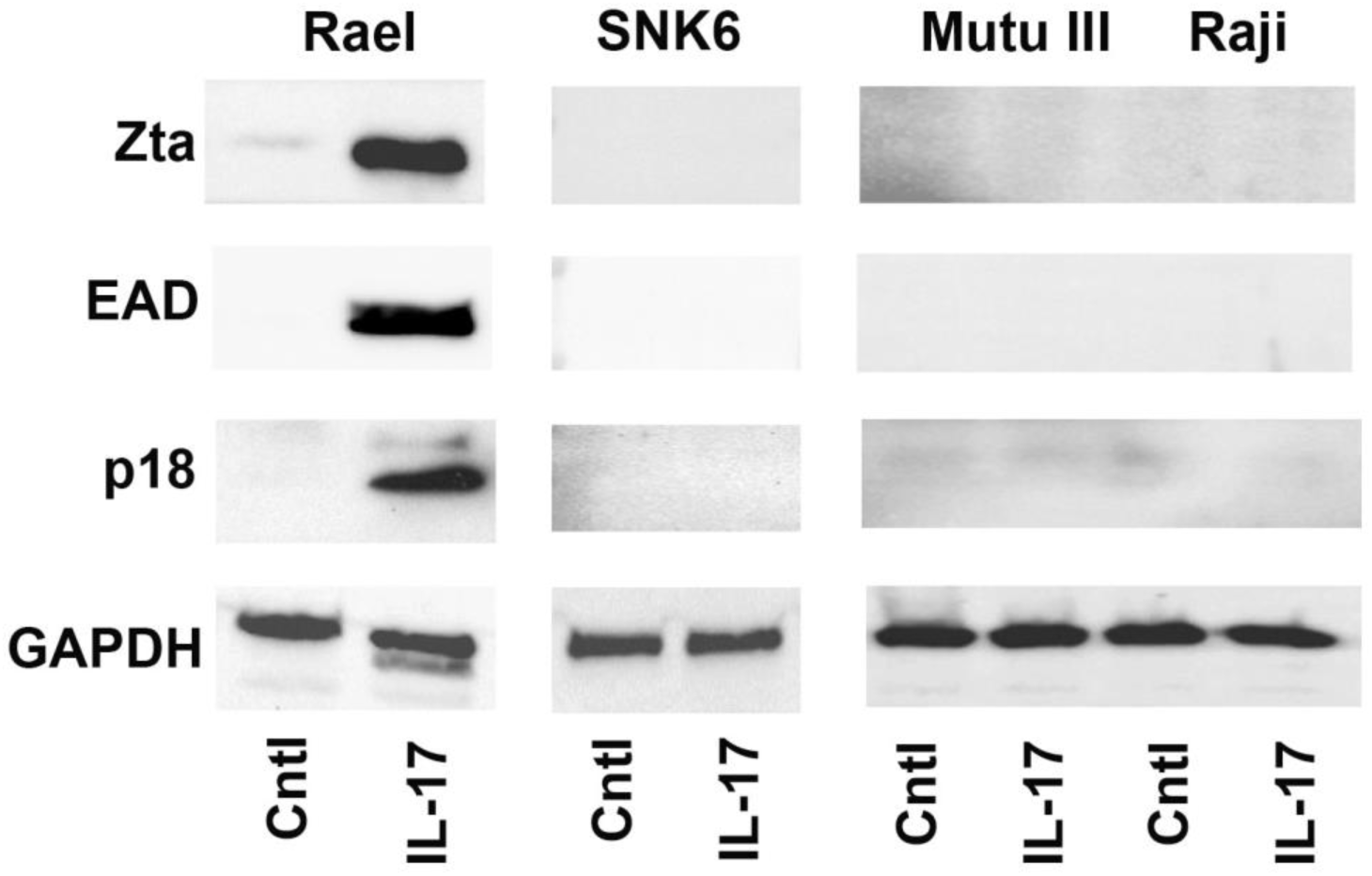
IL-17 preferentially promotes lytic entry in EBV latency I B-cell models. Rael (latency I), SNK6 (latency II), Mutu III (latency III), and Raji (latency III) EBV+ B-cell lines were treated with vehicle or IL-17 (500 ng/ml) for 48 h. Whole-cell lysates were analyzed by immunoblotting for lytic proteins (Zta, EAD, and VCA as available for each line). GAPDH served as a loading control. IL-17 induced lytic protein expression in Rael but not in SNK6, Mutu III or Raji.

## DISCUSSION

Our data show that interleukin-17 (IL-17) alone is sufficient to trigger EBV lytic reactivation and downstream productive events in latently infected human B cells. IL-17 increased the immediate-early regulator BZLF1 (Zta) and early/late transcripts by RT-qPCR and induced lytic proteins (Zta/EAD/p18) by Western blot, establishing concordant transcriptional and protein-level activation of the lytic program. Importantly, culture supernatants from IL-17-treated cells contained significantly higher levels of DNase-resistant extracellular EBV DNA, consistent with encapsidated virions rather than naked DNA released by dying cells. These findings indicate that IL-17 not only initiates lytic entry but also supports productive stages culminating in virion release.

The response followed a clear dose-response. The lytic markers emerged at ∼100 ng/mL and plateaued near 500 ng/mL, which we used to ensure maximal and reproducible activation. Thus, IL-17 defines a cytokine-driven route to EBV reactivation that progresses beyond early gene induction to measurable production of DNase-resistant particles. Neutralization experiments directed at the IL-17A/IL-17R axis are being optimized to provide additional evidence for receptor dependence and to quantify stoichiometric requirements.

Classically, EBV lytic reactivation in B cells has been modeled using B-cell receptor (BCR) cross-linking, phorbol esters/HDAC inhibitors, or tissue-level cues such as hypoxia/HIF-1α, COX-2/PGE₂, and TGF-β, which converge on Zp/Rp via AP-1, CREB/ATF, NF-κB, and chromatin remodeling. These stimuli have been informative yet are often broad or non-physiologic. By contrast, IL-17 is a defined immune cytokine present in inflamed niches that signals through IL-17RA/RC-Act1 (TRAF3IP2) to engage modules already implicated in Zta/Rta control. The demonstration that IL-17 alone elicits both lytic entry and DNase-resistant EBV DNA elevates cytokine signaling from a permissive backdrop to a sufficient upstream trigger of productive reactivation. This positioning invites combinatorial tests (e.g., whether subthreshold IL-17 synergizes with low-dose BCR stimuli, hypoxia, or TGF-β) to map where cytokine inputs intersect with established lytic pathways.

Our transcriptome analyses reinforce this framework. Line-specific Hallmark GSEA identified 13 concordant pathways (FDR < 0.05 in both Akata and Mutu I) dominated by Inflammatory Response, TNFα signaling via NF-κB, Interferon-γ Response, and IL-2/STAT5 signaling, with positive NES in both lines. An orthogonal pooled GO:BP over-representation analysis recovered regulation of B-cell receptor signaling and regulation of B-cell activation, together with cellular response to cytokine stimulus. These pathways support a mechanistic route in which IL-17 engages NF-κB/JNK/ERK/p38 and mRNA-stabilization programs (including IκBζ/NFKBIZ) to lower the threshold for BZLF1/BRLF1 activation. At the receptor level, IL17RA/IL17RC transcripts were readily detectable at baseline, and the adaptor TRAF3IP2/Act1 increased ∼1.5-fold in Akata and ∼10% in Mutu, suggesting cell-line-specific tuning of proximal IL-17 signal strength. Notably, GO terms such as negative regulation of cytokine production, negative regulation of viral process, and ncRNA-mediated gene silencing point to concurrent host feedback/antiviral and chromatin/miRNA modules that could modulate the extent of productive output.

Intriguingly, the gene-level changes indicate IL-17R-mediated NF-κB/MAPK activation and B-cell inflammatory programs. Among the most significantly altered transcripts in Akata were EGR2, EGR1, FCRL5, DOCK10, SCIMP, RGS1, MMP7, and ITGB2. In Mutu, IL-17 up-regulated chemokines (CCL3/CCL3L3/CCL4L1), B-cell activation markers (FCRL5, ITGAX), and a broad interferon-stimulated gene set (IFIT1, IFIT3, RSAD2, IFITM1, ISG15, OAS2, CMPK2, SAMD9L) together with EGR1. These profiles align with the Hallmark GSEA (Inflammatory Response, TNFα/NF-κB, IFN-γ Response, IL-2/STAT5) and the pooled GO:BP enrichment for regulation of B-cell receptor signaling and regulation of B-cell activation. Induction of EGR1/2 (MAPK-responsive immediate-early factors) and SCIMP/RGS1/PTPN6 (B-cell signaling modulators) supports a model in which IL-17R/Act1-TRAF6-TAK1/IKK and MAPKs lower the threshold for BZLF1 transactivation, while the ISG signature in Mutu likely reflects antiviral sensing of IL-17-triggered lytic products. Together, the gene-level changes provide mechanistic granularity beneath the pathway-level results and are concordant with IL-17-driven lytic reactivation.

Our data, together with established signaling maps, support a model in which distinct proximal modules converge on shared distal nodes to license EBV lytic entry (**Fig 6**). In B-cell receptor (BCR) signaling, antigen engagement activates SYK/BTK-PLCγ2-PKCβ, promoting assembly of the CARMA1-BCL10-MALT1 (CBM) complex that recruits TRAF6 and drives K63-linked ubiquitination, thereby activating TAK1-IKK and MAPKs to induce NF-κB (41–43). In IL-17 receptor signaling, Act1/CIKS (TRAF3IP2) couples the receptor to TRAF6, triggering TAK1-IKK and JNK/p38 activation and engaging post-transcriptional programs.

**FIG 6.**
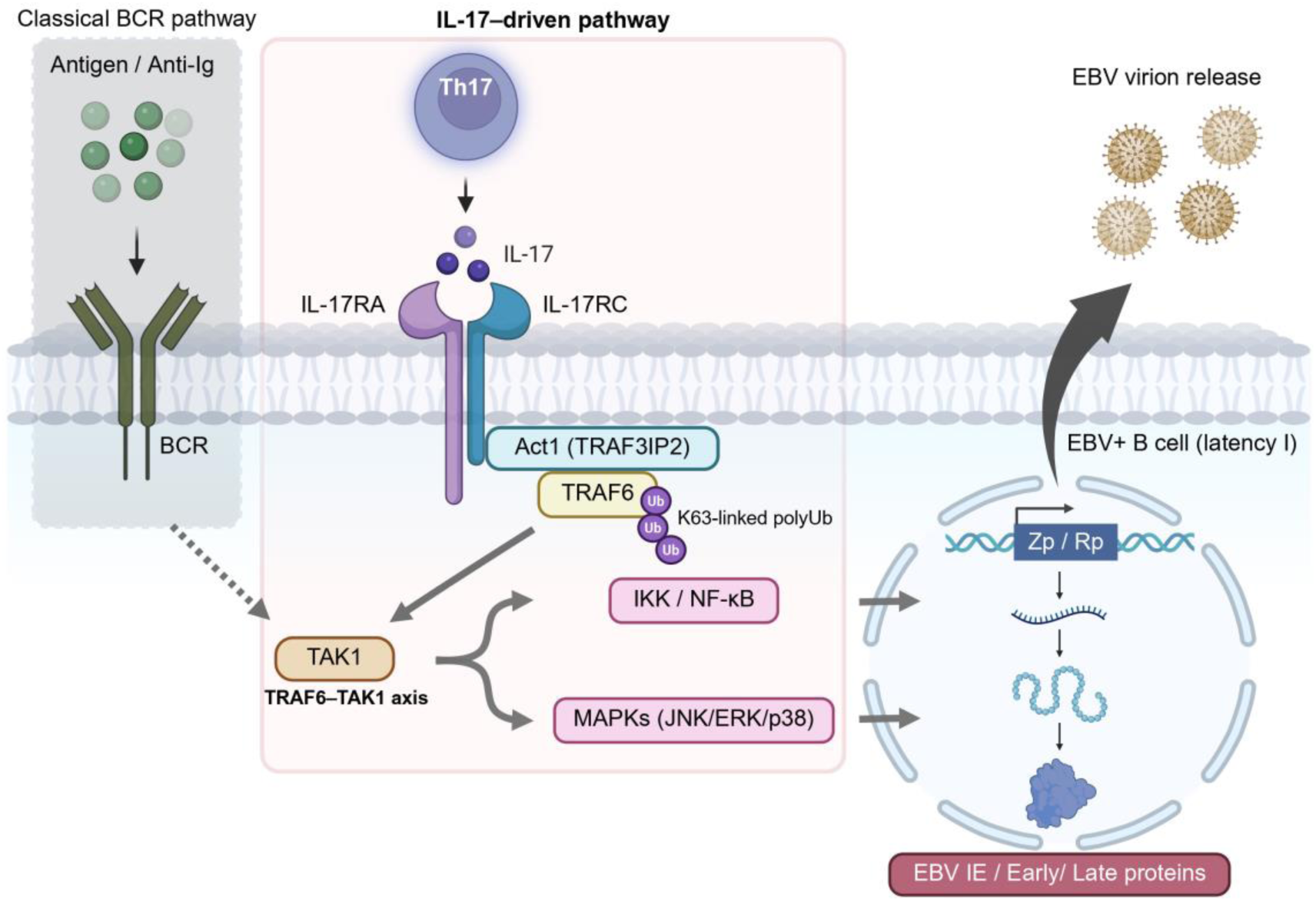
Model schematic: IL-17 signaling and EBV reactivation. IL-17 produced by Th17 cells binds IL-17RA/RC on EBV+ B cells and signals via Act1 (TRAF3IP2) and TRAF6 to activate TAK1, IKK/NF-κB, and MAPKs. In a parallel pathway, B-cell receptor (BCR) engagement activates SYK/BTK-PLCγ2/PKCβ and the CARD11-BCL10-MALT1 (CBM) complex, which also recruits TRAF6 and stimulates the same TAK1-IKK/MAPK nodes. Either pathway alone can therefore drive Zp/Rp (BZLF1/BRLF1) transactivation, leading to EBV lytic gene expression and virion production. Convergence on shared distal effectors predicts that inhibitors of TRAF6-TAK1-IKK/MAPKs will blunt lytic reactivation induced by both IL-17 and BCR, while perturbation of pathway-specific nodes will differentially affect the two inputs. Created in BioRender. Dochi, H. (2025) https://BioRender.com/xhy86xa.

Pharmacologic and genetic analyses place TAK1 upstream of IL-17-dependent MAPKs (44–46). Although the proximal adaptors differ (Act1-centered for IL-17R vs SYK/BTK/PLCγ2/PKCβ/CBM for BCR), both pathways converge on TRAF6-TAK1-IKK and MAPKs, the same distal effectors long implicated in Zp/Rp transactivation via AP-1/ATF and stress-kinase inputs (14, 17, 47). This biochemical convergence provides a mechanistic rationale for why IL-17 can phenocopy or potentiate BCR-like outputs that lower the threshold for the EBV immediate-early switch, and it yields testable predictions: shared-node blockade (e.g., TAK1 or IKK inhibition) should blunt both IL-17- and BCR-induced lytic readouts, whereas pathway-specific perturbations (e.g., Act1/CIKS disruption versus BTK/CARD11/MALT1 inhibition) should differentially attenuate the two inputs.

These findings have pathobiologic and translational implications. Many EBV-associated settings display Th17-skewed inflammation with elevated IL-17. Our results suggest that IL-17-rich microenvironments may promote episodic, productive reactivation, increasing local virion burden and antigen exposure, with consequences for immune activation, bystander damage, and tumor-immune dynamics. A feed-forward model is therefore plausible. EBV products can stimulate IL-17 responses, and IL-17 can reciprocally augment EBV reactivation, establishing an inflammation-reactivation loop. This yields testable predictions.

Thus, IL-17/IL-17R blockade may reduce EBV reactivation markers in IL-17-high contexts, while conditions that amplify Th17 responses may raise the lytic burden. Because we detect DNase-resistant extracellular EBV DNA after IL-17 exposure, future clinical studies could correlate tissue or plasma IL-17 with cell-free EBV DNA as a practical readout of reactivation pressure *in vivo*.

## CONCLUSIONS

Our data identify IL-17 as a sufficient cytokine trigger of Epstein-Barr virus (EBV) lytic reactivation in human B cells. The response was observed across multiple EBV+ B-cell lines with latency program I, indicating that IL-17-responsive reactivation is not confined to a single model. These findings place IL-17 signaling alongside established lytic inducers and provide a biologically plausible route by which Th17-skewed inflammation could increase EBV reactivation. Collectively, our results establish the IL-17 pathway as a directly actionable regulator of the EBV latency-lytic switch and provide a framework for mechanistic dissection and translational testing.

## ACKNOWLEDGEMENTS

Research reported in this publication was supported by the National Cancer Institute of the National Institutes of Health under Award Number R01CA261258 to ZL. The content is solely the responsibility of the authors and does not necessarily represent the official views of the National Institutes of Health. This work was also supported by a U.S.-Japan Cooperative Medical Sciences Program Collaborative Award from the National Institute of Allergy and Infectious Diseases and CRDF Global (grant number DAA3-19-65602-1), a Louisiana Cancer Research Center pilot grant, a Carol Lavin Bernick faculty grant, and a Ladies Leukemia League research grant to ZL.

